# Loss of active neurogenesis in the adult shark retina

**DOI:** 10.1101/2020.11.19.389973

**Authors:** Ismael Hernández-Núñez, Diego Robledo, Hélène Mayeur, Sylvie Mazan, Laura Sánchez, Fátima Adrio, Antón Barreiro-Iglesias, Eva Candal

**Author notes:** Equal contributors (Antón Barreiro-Iglesias, Eva Candal). Correspondence: Eva Candal.

## Abstract

Neurogenesis is the process by which progenitor cells generate new neurons. As development progresses neurogenesis becomes restricted to concrete neurogenic niches, where it persists during postnatal life. The retina of teleost fishes is thought to proliferate and produce new cells throughout life. Whether this capacity may be an ancestral characteristic of jawed vertebrates, shared with chondrichthyans, which diverged from osteichthyans prior to the gnathostome radiation is completely unknown. Previous work from our group revealed that the juvenile retina of the catshark Scyliorhinus canicula shows active proliferation and neurogenesis. Here, we compared the morphology and proliferative status of the retina between catshark juveniles and adults. Histological analyses revealed an important reduction in the size of the peripheral retina (where progenitor cells are mainly located), an increase in the thickness of the plexiform layers and a decrease in the thickness of the inner nuclear layer in adults. Contrary to what has been reported in teleost fish, we did not observe active mitotic activity in the catshark retina after sexual maturation, suggesting that there is no significant proliferation and neurogenesis in adult specimens. Based on these results, we carried out RNA-Sequencing (RNA-Seq) analyses comparing the retinal transcriptome of juveniles and adults, which revealed a statistically significant decrease in the expression of many genes involved in cell proliferation and neurogenesis in adult catsharks. Our RNA-Seq data provides an excellent resource to identify new signaling pathways controlling neurogenesis in the vertebrate retina.

## 1. Introduction

Neurogenesis is the process by which neural progenitor cells generate new neurons. In vertebrates, adult neurogenesis only occurs in specific regions of the central nervous system (CNS) called neurogenic niches. Teleost fish display the most prominent and widespread adult neurogenesis throughout the CNS compared with any other vertebrate studied so far (Ganz and Brand, 2016). From anamniotes to amniotes, constitutive neurogenesis in adults become successively restricted to fewer neurogenic niches (Grandel and Brand, 2012). Actually, the occurrence of adult neurogenesis in certain neurogenic niches of mammals, including humans, is still controversial (Boldrini et al., 2018; Paredes et al., 2018; Sorrells et al., 2018; Tobin et al., 2019).

The fact that the retina of teleost fishes shows active proliferation and neurogenic activity throughout life (goldfish: Johns, 1977; rainbow trout: Julian et al., 1998; zebrafish: Marcus et al., 1999) together with its cytoarchitectonic simplicity and amenability to manipulation provided the basis for the use of the teleost retina as a model for the study of adult neurogenesis in vertebrates. Two main niches of constitutive neurogenesis have been described in the retina of adult teleosts: the ciliary marginal zone (CMZ), which is a continuously growing circumferential ring of cells located in the peripheral part of the retina (Johns, 1977; Raymond et al., 2006; Wan et al., 2016), and Müller glial cells located in the central (mature) retina which can act as progenitor cells in constitutive and regenerative conditions (Fausett and Goldman, 2006; Bernardos et al., 2007; Hamon et al., 2016, 2017; Jorstad et al., 2017; Hamon et al., 2019).

While the distribution of adult neurogenic niches shows an outstanding conservation among teleost species (Ganz and Brand, 2016), the distribution of neurogenic niches within the retina of different vertebrates might not so well conserved. Actually, the retina of adult lampreys (a jawless vertebrate) shows no proliferative activity (Villar-Cheda et al., 2008), and some vertebrate groups show constitutive neurogenesis in the adult retina from niches other than the CMZ and Müller glial cells (reviewed in Amato et al., 2004; Moshiri et al., 2004). Besides, in fishes, data on adult neurogenesis in the retina come only from studies in modern teleost models (e.g. zebrafish or goldfish) but there is no data on adult neurogenesis on more basal teleosts, other ray-finned fishes (e.g. chondrosteans) or in chondrichthyans (cartilaginous fishes, the oldest extant jawed vertebrates). Therefore, a better understanding of adult neurogenesis in a representative sampling of vertebrates, including groups which diverged early from the osteichthyan lineage, will be essential to gain knowledge on the evolution of adult neurogenesis in the vertebrate retina.

As the sister group of osteichthyans, cartilaginous fishes occupy a key phylogenetic position to infer ancestral states fixed prior to the gnathostome radiation or identify lineage-specific diversifications in the major osteichthyan taxa. Our research group has thoroughly studied the development of the retina of an elasmobranch fish, the catshark Scyliorhinus canicula. The developing retina of S. canicula shows a high proliferative activity that decreases throughout development, although it is still observed in juveniles (Ferreiro-Galve et al., 2010). An interesting feature of the retina of S. canicula is that it presents a region adjacent to the CMZ, called the transition zone (TZ), which also contains progenitor cells (Ferreiro-Galve et al., 2010; 2012; Sánchez-Farías and Candal, 2015, 2016; Hernández-Núñez et al., unpublished). Different types of progenitor cells have been identified in the CMZ and TZ both in the developing and in the postnatal retina of S. canicula based on the expression of progenitor cell markers like proliferating cell nuclear antigen (PCNA), phosphohistone H3 (pH3), glial fibrillary acidic protein (GFAP), Sox2, or brain lipid-binding protein (Ferreiro-Galve et al., 2010; 2012; Sánchez-Farías and Candal 2015, 2016, Hernández-Núñez et al., unpublished). However, whether this neurogenic activity is maintained in the adult retina has not been addressed so far.

Here, we aimed to gain knowledge about the evolution of adult neurogenesis in the vertebrate retina by exploring this process in cartilaginous fishes with the catshark Scyliorhinus canicula as reference. We characterized changes in the morphology and neurogenic ability of the retina of S. canicula from postnatal development to adulthood using classical histological staining and by analysing the expression of proliferation markers. Additionally, we carried out an Illumina RNA-sequencing (RNA-Seq) study comparing the retinal transcriptomes of juveniles and adults to detect changes in the expression of genes involved in cell proliferation and neurogenesis, and to identify new genes and signaling pathways responsible for the neurogenic process in the retina of vertebrates.

## 2. Material and Methods

### 2.1. Animals

Juvenile (n = 15; 10 to 13 cm long) and adult (n = 14; 45 to 50 cm long) specimens of S. canicula were kindly provided by the aquarium Acuario do Grove (O Grove, Spain) and kept in seawater tanks under standard conditions of temperature (15-16 °C), pH (7.5-8.5) and salinity (35 g/L). All experimental procedures were performed following the guidelines established by the European Union and the Spanish government for animal experimentation and were approved by the Bioethics Committee of the University of Santiago de Compostela.

### 2.2. Tissue preparation for histology

Animals were deeply anesthetized with 0.5 % tricaine methanesulfonate (MS-222, Sigma, St. Louis, MO) in seawater and then perfused intracardially with elasmobranch Ringer’s solution (see Ferreiro-Galve et al., 2008) followed by perfusion with 4% paraformaldehyde (PFA) in 0.1M elasmobranch phosphate buffer (pH 7.4) containing 1.75% urea. The eyes were removed and postfixed in 4% PFA for 2 days at 4 °C. After rinsing in phosphate buffer saline (PBS), the eyes were cryoprotected with 30 % sucrose in PBS, embedded in Neg-50TM (Thermo Scientific, Kalamazoo, MI) and frozen with liquid nitrogen-cooled isopentane. Transverse sections (18 μm thick) were obtained on a cryostat and mounted on Superfrost Plus slides (Menzel-Glasser, Madison, WI).

### 2.3. Haematoxylin-eosin staining

Some retinal sections were stained with haematoxylin-eosin following standard protocols. Briefly, cryostat sections were dried at room temperature, rinsed in 0.05 M Tris-buffered (pH 7.4) saline (TBS) for 10 min and stained with haematoxylin solution for 10 min. Sections were subsequently rinsed in tap water until removal of the excess of haematoxylin, in distilled water for 10 min and then stained with eosin for 2 min. Finally, the sections were dehydrated and mounted in DPX mounting medium (Scharlau, Sentmenat, Spain).

### 2.4. Immunofluorescence

Sections were first pre-treated with 0.01 M citrate buffer pH 6.0 for 30 min at 90 °C for heat-induced epitope retrieval, allowed to cool for 20 min at room temperature (RT) and rinsed in TBS for 5 min. Then, sections were incubated with different combinations of primary antibodies [GFAP/glutamine synthase (GS) or PCNA/pH3; Supplementary Table 1] overnight at RT. Sections were rinsed three times in TBS for 10 min each, and incubated in a combination of fluorescent dye-labelled secondary antibodies (Supplementary Table 1) for 1 h at RT. All antibody dilutions were made in TBS containing 15 % normal goat serum (Millipore, Billerica, MA), 0.2 % Triton X-100 (Sigma) and 2 % BSA (Sigma). Sections were then rinsed three times in TBS for 10 min each and in distilled water for 30 min, allowed to dry for 30 min at 37 °C and mounted in MOWIOL 4-88 Reagent (Calbiochem, Darmstadt, Germany).

### 2.5. Specificity of antibodies

The anti-GFAP and the anti-GS antibodies have been previously used in the retina of the juvenile catshark as markers of early and late glial cells, respectively (Sanchez-Farías and Candal, 2016) and their specificity has been tested by western blot in brain protein extracts of adult catsharks (Docampo-Seara et al., 2019). PCNA is present in proliferating cells and although its expression is stronger during the S phase, it persists along the entire cell cycle excepting the mitotic period (Zerjatke et al., 2017). The anti-PCNA antibody has been previously used to label progenitor cells in the brain and retina of S. canicula (i.e. Quintana-Urzainqui et al., 2014 Sanchez-Farías and Candal, 2015). The anti-pH3 antibody has been also widely used in the brain and retina of S. canicula as a marker of mitotic cells (Ferreiro-Galve et al., 2010; Quintana-Urzainqui et al., 2014; Docampo-Seara et al., 2019).

### 2.6. Riboprobe synthesis and *in situ* hybridization

*In situ* hybridization (ISH) experiments were carried out to study the expression of S. canicula Sox2 (*ScSox2*) transcripts using the probe described in Lagadec et al. (2018). Sense and antisense digoxigenin-UTP-labelled *ScSox2* riboprobes were synthesized by in vitro transcription, after PCR amplification, and using the T7 or SP6 RNA polimerases (NZYTech, Lisbon, Portugal). ISH was performed on cryostat sections (18 μm) of juvenile and adult retinas as previously described (Coolen et al., 2007). Briefly, sections were permeabilized with proteinase K, hybridized with sense or antisense probes overnight at 65 °C and incubated with the alkaline phosphatase-coupled sheep anti-digoxigenin antibody (1:2000, Roche Applied Science, Manheim, Germany) overnight at 4 °C. The colour reaction was performed using BM-Purple (Roche).

Finally, sections were dehydrated and mounted in DPX. Control sense probes did not produce any detectable signal.

### 2.7. Image acquisition

Images of fluorescent labelled sections were taken with a Leica TCS-SP2 confocal microscope with a combination of blue and green excitation lasers. Confocal optical sections were taken at steps of 1 μm along the z-axis. Collapsed images were obtained with the LITE software (Leica). Brightfield images were obtained with an Olympus BX51 microscope equipped with an Olympus DP71 camera. Contrast and brightness were minimally adjusted using Adobe Photoshop CS4 (Adobe, San Jose, CA).

### 2.8. Cell quantifications and statistical analyses

We quantified the number of mitotic [pH3 positive (pH3+) cells] in the peripheral retina (including the CMZ and TZ) and proliferative cells (PCNA+ cells) in the central retina of juveniles (n = 4 retinas from 4 individuals) and adults (n = 4 retinas from 4 individuals). The number of pH3+ cells and PCNA+ cells were manually counted under the microscope in one out of each four consecutive retinal sections (18 μm). The limit between the peripheral and the central retina was established based on their morphological differences like the characteristic layered structure of the central retina. Then, we calculated the mean number of cells per section for each retina. Statistical analyses were performed with Prism 8 (GraphPad software, La Jolla, CA). Normality of the data was determined with the Shapiro Wilk test. A Mann Whitney test (pH3+ cells) and a Student’s (unpaired) t-test (PCNA+ cells) were used to determine significant differences (p < 0.05) between juveniles and adults.

### 2.9. RNA isolation and sequencing

The retinas of juveniles (n = 6 retinas from 5 animals) and adults (n = 5 retinas from 4 animals) were dissected out and put in RNAlater (Ambion Inc., Austin, TX, USA). RNA extraction was performed using the RNeasy mini kit (Qiagen, Hilden, Germany) with DNase treatment following the manufacturer’s instructions. Isolated RNAs were eluted in nuclease free water. RNA quality and quantity were evaluated in a Bioanalyzer (Bonsai Technologies, Madrid, Spain) and in a NanoDrop^®^ ND-1000 spectrophotometer (NanoDrop^®^ Technologies Inc., Wilmington, DE, USA). Thereafter, the Illumina Truseq mRNA stranded RNA-Seq Library Prep Kit protocol was followed. Libraries were checked for quality and quantified using the Bioanalyzer 2100 (Agilent, Santa Clara, CA, USA) (all samples had RIN values higher than 7), before being sequenced on one S1 lane of the Illumina NovaSeq instrument using 150 base paired-end sequencing at Edinburgh Genomics (UK). Raw reads have been deposited in NCBI’s Sequence Read Archive (SRA) under BioProject accession number PRJNA668789.

The quality of the sequencing output was assessed using FastQC v.0.11.5 (http://www.bioinformatics.babraham.ac.uk/projects/fastqc/). Quality filtering and removal of residual adaptor sequences was conducted on read pairs using Fastp v.0.20.0 (Chen et al., 2018). Illumina specific adaptors were clipped from the reads and leading and trailing bases with a Phred score less than 20 were removed; only reads where both pairs were longer than 36 bp post-filtering were retained. Filtered reads were mapped to a catshark transcriptomic reference obtained by a clustering approach (Mayeur & Mazan, to be published elsewhere) and transcript abundance was quantified using Kallisto v0.46.1 (Bray et al., 2016).

Differential expression (DE) analyses were performed using R v.3.6.2 (https://www.r-project.org/). Gene count data were used to estimate differential gene expression using the Bioconductor package DESeq2 v.3.4 (Love et al., 2014). Briefly, size factors were calculated for each sample using the ‘median of ratios’ method and count data was normalized to account for differences in library depth. Next, gene-wise dispersion estimates were fitted to the mean intensity using a parametric model and reduced towards the expected dispersion values. Finally, a negative binomial model was fitted for each gene and the significance of the coefficients was assessed using the Wald test. The Benjamini-Hochberg false discovery rate (FDR) multiple test correction was applied, and genes with FDR < 0.01, normalized mean read counts > 50 and absolute log2 fold change values (FC) > 1 were considered differentially expressed. Gene Ontology (GO) and Kyoto Encyclopedia of Genes and Genomes (KEGG) enrichment were performed using the DAVID bioinformatics resource Functional Annotation tool (Huang et al., 2009), using the shark transcriptome as background.

## 3. Results

### 3.1. Comparison of juvenile and adult retinal morphology

As in lampreys and jawed vertebrates, the catshark retina is composed of three nuclear layers [the outer nuclear layer (ONL), the inner nuclear layer (INL) and the ganglion cell layer (GCL)] and two plexiform layers [the outer plexiform layer (OPL) and the inner nuclear layer (INL)]. The adult retina increases in size respect to the juvenile retina. As previously reported (Ferreiro-Galve et al., 2010), the peripheral retina of juveniles showed a non-layered CMZ with a neuroepithelial organization and a TZ where the IPL separates the GCL from an unlayered region (Fig. 1A). In the central retina, we observed the typical layered organization (Fig. 1B). In adults, the peripheral retina showed the same zones observed in juveniles but with an important reduction in size and extension in comparison to the juvenile retina and also relative to the total size of the adult retina (Fig. 1C). The adult central retina also showed the typical layered organization, but we observed a decrease in the thickness of the INL and an increase in the thickness of the two plexiform layers (Fig. 1D).

**Figure 1.**
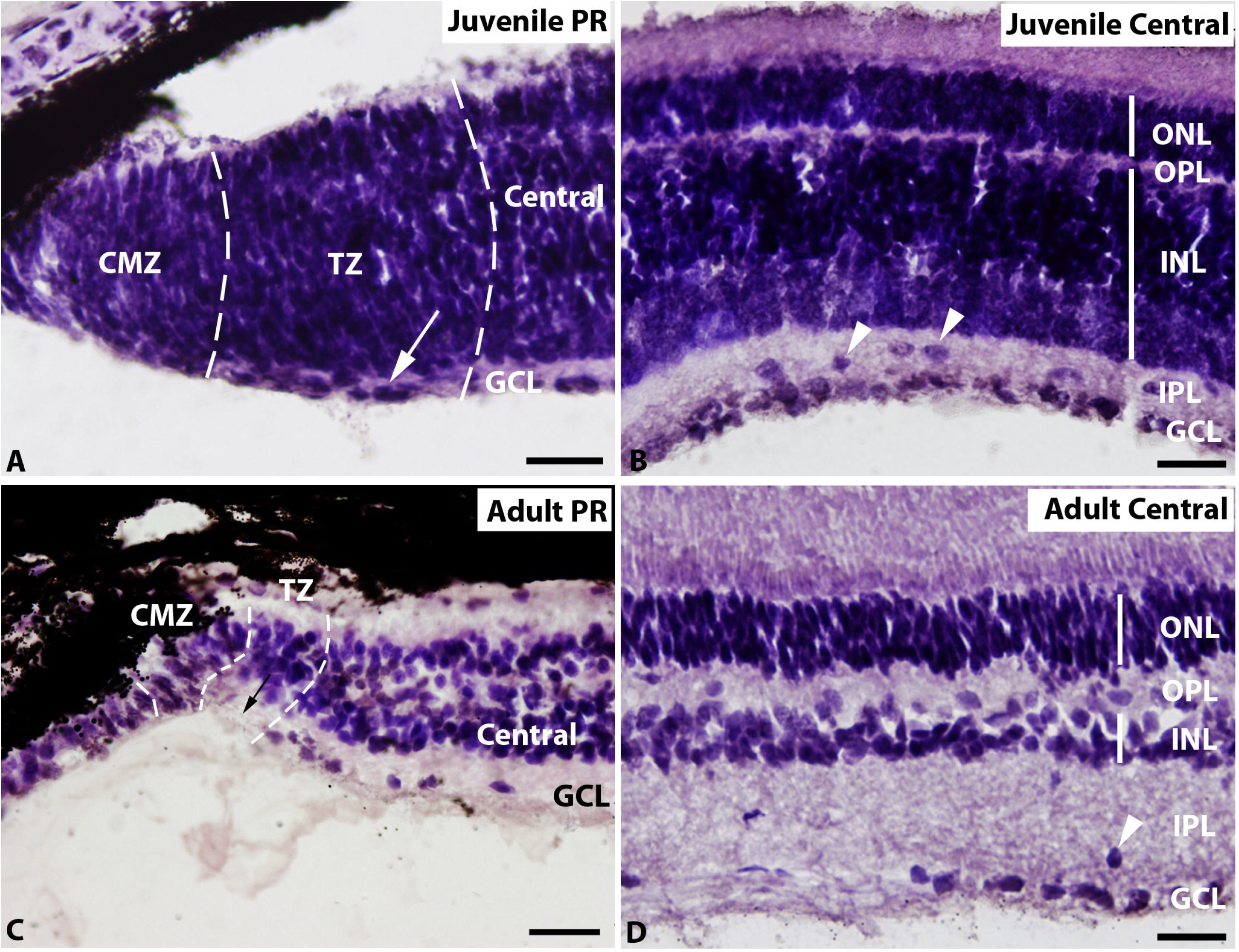
Haematoxylin-eosin stained transverse sections of the retina of juvenile (A, B) and adult (C, D) specimens of S. canicula. (A) Juvenile peripheral retina showing an unlayered CMZ and a TZ with an IPL (arrow) separating the GCL from an unlayered region. (B) Juvenile central retina showing all the characteristic layers: GCL, IPL, INL, OPL, ONL and displaced cells in the IPL (arrowheads). (C) The adult peripheral retina showed a reduction in size and extension in comparison to the juvenile retina. Note also the increase in the thickness of the IPL in the TZ and central retina (arrow). (D) The adult central retina showed an evident decrease in the thickness of the INL and an increase in the thickness of the OPL and IPL. Displaced cells were observed in the IPL (arrowheads). Scale bars: 100 μm (A); 50 μm (B, C, D). Abbreviations: CMZ: ciliary marginal zone; GCL: ganglion cell layer; INL: inner nuclear layer; IPL: inner plexiform layer; ONL: outer nuclear layer; OPL: outer plexiform layer; PR: peripheral retina; TZ: transition zone.

### 3.2. Defining the peripheral retina

To better define the borders of the peripheral retina, where the main retinal neurogenic niche is located, we studied the expression pattern of two glial cell markers (GFAP and GS) and the stem cell marker Sox2.

In the juvenile retina, GFAP expression was observed in processes of progenitor cells in the CMZ and TZ (Fig. 2A; Sánchez-Farías and Candal, 2016) and in mature Müller glial cells in the central retina (Figs. 2A, B; Sánchez-Farías and Candal, 2016). However, GS expression was restricted to processes and cell bodies of mature Müller glial cells in the central retina (Fig. 2B), which facilitates delimiting the boundary between the peripheral and central retinas (Fig. 2A).

**Figure 2.**
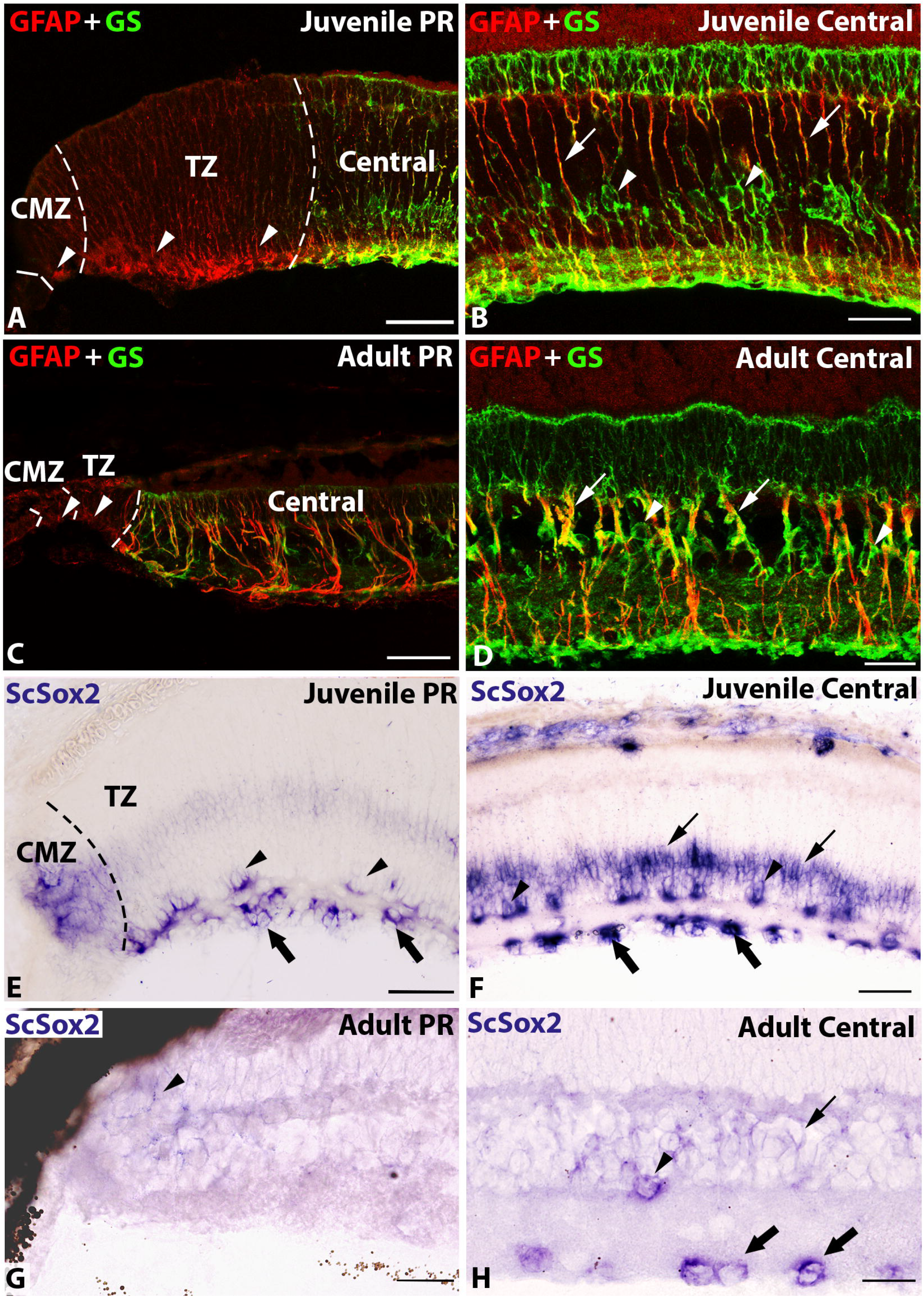
Double labelling with GFAP and GS (A-D) and *in situ* hybridization of *ScSox2* (E-H) in transverse sections of the retina of juvenile and adult specimens of S. canicula. (A) In the juvenile retina, GFAP was expressed in processes of progenitor cells in the CMZ and TZ (arrowheads) while GS expression was not observed in this region. (B) In the juvenile central retina, coexpression of GFAP and GS was observed in processes (arrows) and GS in cell bodies (arrowheads) of Müller glia. (C) In the adult peripheral retina, GFAP+ putative progenitor cells are highly restricted to the retinal margin (arrowheads). (D) In the adult central retina, GFAP+ and GS+ Müller glia processes (arrows) and GS+ Müller glia cell bodies (arrowheads) were observed as in juveniles. (E) *ScSox2* expression was observed in the CMZ and basal region of the TZ of the juvenile retina. Note the presence of amacrine (arrowheads) and ganglion (thick arrows) cells in the TZ. (F) *ScSox2* expression was observed in amacrine (arrowheads), ganglion (thick arrows) and Müller glia (arrows) cells of the juvenile retina. (G) *ScSox2* expression was much weaker (arrowhead) in the adult peripheral retina than in juveniles. (H) In the adult central retina, the pattern of expression of *ScSox2* was similar to juveniles but weaker. Note the presence of *ScSox2* positive amacrine (arrowheads) and ganglion (thick arrows) cells. Scale bars: 100 μm (A, C, E, G); 25 μm (B, D, F, H). Abbreviations: CMZ: ciliary marginal zone; PR: peripheral retina; TZ: transition zone.

In adults (Figs. 2C, D), the expression patterns of GFAP and GS were similar to those observed in juveniles (Figs. 2A, B), and also allowed for a clear identification of peripheral and central regions of the retina (Figs. 2C, D). The size of the peripheral retina containing GFAP+ putative progenitors in adults was highly reduced with respect to juveniles and relative to the size of the entire retina (Fig. 2C).

Sox2 is known to be expressed in progenitor cells that keep their stem cells properties in adults (DeOliveira-Mello et al., 2019). *ScSox2 in situ* labelling allowed us to clearly establish the boundary between the CMZ and TZ within the peripheral retina in the juvenile retina (Figs. 2E). An intense expression of *ScSox2* was observed in the CMZ and the basal region of the TZ (Fig. 2E), and in amacrine, ganglion and Müller glia cells (Fig. 2F) in the central retina of juveniles. The same pattern of expression was observed in adults (Figs. 2G, H), although *ScSox2* expression in the adult retina (Fig. 2G, H) clearly decreased with respect to that observed in juveniles (Fig. 2E, F).

### 3.3. Proliferation and mitotic activity

It is well known that the proliferative potential of the retina decreases during development in vertebrates; nevertheless, cell proliferation is still observed in adult teleost fish (see introduction). Here, we compared the proliferative status and mitotic activity of juvenile and adult catshark retinas using PCNA and pH3 double immunolabeling. As previously reported (Ferreiro-Galve et al., 2010), in the peripheral retina of juveniles abundant PCNA+ cells were found throughout the CMZ and the outer part of the TZ, but not in the prospective GCL (Fig. 3A). pH3 positive (pH3+) cells were restricted to the ventricular (apical) region of the peripheral retina (Fig. 3A). In the central retina of juveniles, scattered PCNA+ cells were found in the INL (Fig. 3B), bordering the horizontal cell layer (HCL; not shown) and in the GCL/IPL (Fig. 3B). No pH3+ cells were found in the central retina of juveniles (not shown).

**Figure 3.**
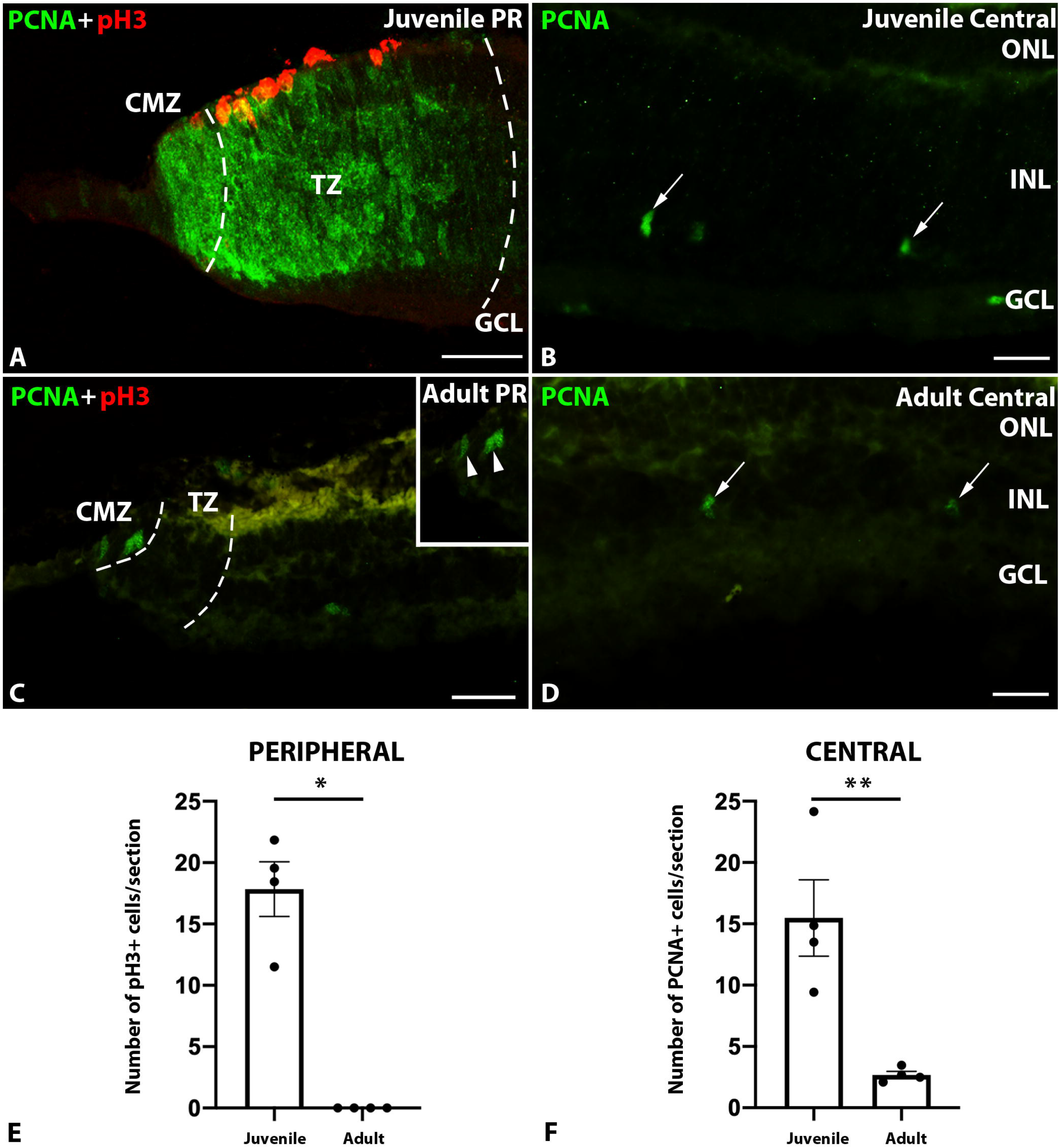
Double labelling with PCNA and pH3 in transverse sections of the retina of juvenile and adult specimens of S. canicula (A-D) and graphical representations of the average number of mitotic (E) and proliferative cells (F) per section in both stages. (A) Abundant PCNA+ cells were observed in the CMZ and the TZ of juveniles. Note the absence of PCNA+ cells in the GCL. Mitotic cells (pH3+) were only found in the ventricular (apical) region. (B) Scattered PCNA+ cells were found in the central retina of juveniles (arrows). (C) In adults, only a few PCNA+ cells were observed in the apical region of the CMZ and TZ (arrowheads in inset), while no pH3+ cells were found in this region. (D) Occasional PCNA+ cells were found in the adult central retina (arrows). (E) Graph showing the average number of mitotic cells (pH3+) per section in the peripheral retina of juveniles and adults revealing statistically significant differences in the number of mitotic cells (p-value = 0.0286; *). (F) Graph showing the average number of proliferating cells (PCNA+) per section in the central retina of juveniles and adults revealing statistically significant differences in the number of proliferating cells (p-value = 0,0064; **). Each dot in the graphs (E, F) represents one retina. Scale bars: 50 μm (A); 25 μm (B, D); 100 μm (C). Abbreviations: CMZ: ciliary marginal zone; GCL: ganglion cell layer; INL: inner nuclear layer; IPL: inner plexiform layer; ONL: outer nuclear layer; OPL: outer plexiform layer; PR: peripheral retina; TZ: transition zone.

In the peripheral retina of adults only occasional PCNA+ cells were observed, most of them being restricted to the retinal margin (Fig. 3C). Interestingly, no pH3+ cells were observed in this region (Fig. 3C). Only occasional PCNA+ cells were observed in the central retina of adults (Fig. 3D). As in juveniles, no pH3+ cells were found in this region (not shown).

Quantifications of mitotic (pH3+) and proliferating (PCNA+) cells revealed significant differences in the proliferative capacity of juvenile and adult retinas. There was a statistically significant decrease in the number of pH3+ cells (p = 0.0286) in the peripheral retina of adults (Fig. 3E). We also observed a statistically significant decrease in the number of PCNA+ cells in the central retina of adults (p = 0.0064, Fig. 3F). The pH3 results reveal a dramatic loss of mitotic activity in adult retinas.

### 3.4. RNA-Sequencing

Our results revealed important differences between the juvenile and the adult retina mainly regarding the extension and distribution of progenitor cells in the peripheral retina and the huge decrease of proliferative activity after sexual maturation, with a complete lack of mitotic activity in the retina of adult specimens.

In order to identify the genes and signaling pathways responsible for the high proliferative activity of the juvenile retina compared to the quiescence observed in adults, we performed an Illumina RNA-Seq analysis comparing the retinal transcriptomes of juveniles and adults. The juvenile and adult retinal transcriptomes showed marked differences and clustered clearly apart in the principal component analysis (Fig. 4A). The first principal component, separating the juvenile and adult samples, explained a very large proportion of the variance (58.3%). Juvenile retinas formed a tight cluster, whereas adult retinas showed more variation, which might be attributed to higher differences in age of the adult individuals. A total of 6,359 differentially expressed genes were detected (Fig. 4B; Supplementary File 1). Of these, 4,203 genes showed decreased expression and 2,156 showed increased expression in the adult retina (Supplementary File 1). GO term and KEGG pathway enrichment analyses showed that many of the significantly enriched pathways are related to cell division and cell cycle arrest (GO terms: “mitotic nuclear division”, “mitotic cytokinesis”, “regulation of exit from mitosis” and “cell cycle arrest”), neuronal plasticity (GO terms: “regulation of neuronal synaptic plasticity” and “dendrite morphogenesis”) or other developmental pathways (GO terms: “regulation of establishment of cell polarity”, “photoreceptor cell maintenance” and “regulation of cellular senescence”, Supplementary File 2). These results were in good agreement with the differences observed regarding changes in proliferative activity between juvenile and adult retinas (see above).

**Figure 4.**
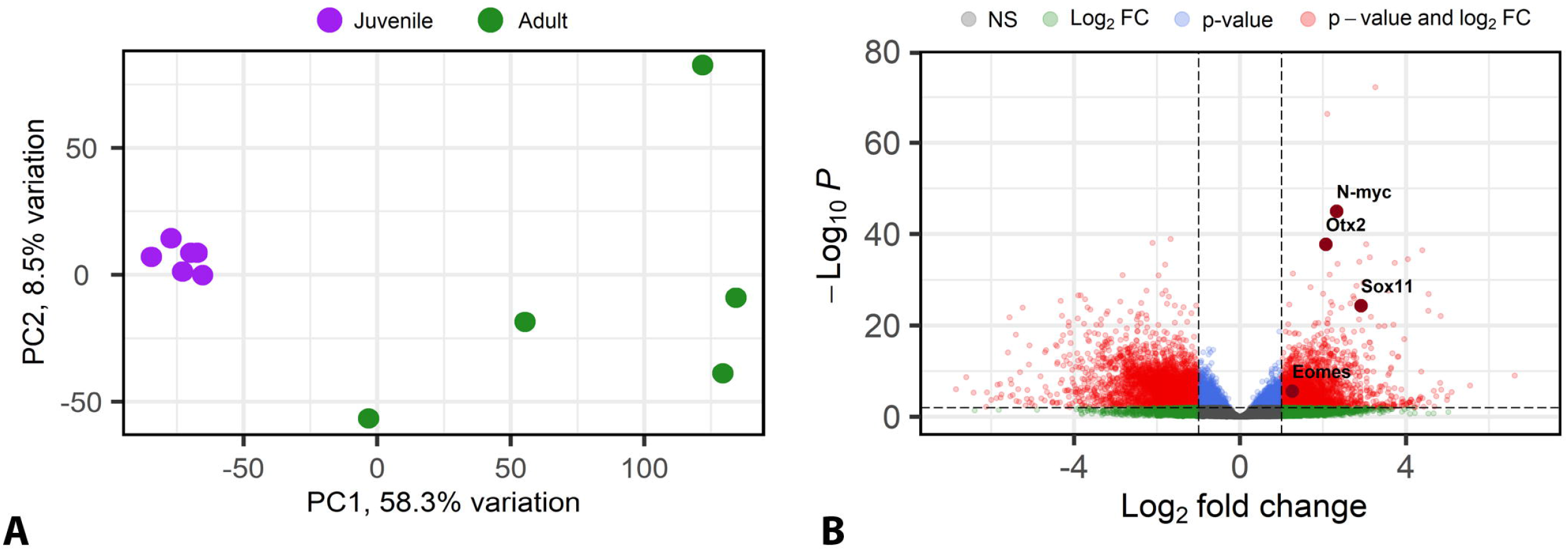
RNA-Seq analysis of the retinal transcriptome (A-B). (A) Principal component analysis showing the juvenile (purple dots) and adult (green dots) samples clustered according to their gene expression. Each dot represents one retina. (B) Volcano plot displaying the results of the RNA-seq analysis (statistical significance vs magnitude of change) and highlighting significant genes. Each dot represents 1 gene. 4 of the genes that showed decreased expression in the adult retina (Eomes, Otx2, N-myc and Sox11) are highlighted. A positive fold change indicates a higher expression in the juvenile retina. Abbreviations: FC: fold change; NS: not significant; PC: principal component.

Genes showing decreased expression in the adult retina are good candidates to be pro-neurogenic and responsible for the higher proliferative activity of the juvenile retina. Actually, many of the genes that showed decreased expression in the adult retina [e.g. several cyclins (e.g. CCNA2, or CCND2), Ki67, N-myc, Otx2, Foxo1, Olig2, Eomes (Tbr2) or Sox family factors like Sox11 (Fig. 4B; Supplementary File 1)] are known to be important players in the maintenance of neural stem cell properties, cell proliferation, cell cycle progression or neurogenesis. Furthermore, genes belongings to signaling pathways known to play major roles in nervous system development (e.g. Shh, Wnt, Notch or Slit-Robo signaling pathways) also showed differential expression (Supplementary Table 2). These results were in good agreement with the histological observations (see above) and revealed that these signaling pathways might play a key role in retinal neurogenesis in sharks.

## 4. Discussion

The teleost retina shows continual growth throughout life (Marcus et al., 1999; Raymond et al., 2006; Wan et al., 2016). In sharks, the eyes also show continuous growth, with a close relationship between eye diameter and body length (Litherland et al., 2009; Collin, 2018). The extension and organization of the peripheral retina observed in juveniles (present results; Ferreiro-Galve et al., 2010) was very similar to that previously described in late embryos (stages 33-34 as defined by Ballard et al., 1993) (Ferreiro-Galve et al., 2010), while in adults it showed a reduction in size compared to juveniles and also relative to the entire size of the adult retina. This decrease in the extension of the peripheral retina has been also observed during early retinal development in S. canicula (Ferreiro-Galve et al., 2010) and in reptile (Eymann et al., 2019) embryos. The organization of the central retina in juveniles and adults was similar to that described in stage 33-34 embryos (Ferreiro-Galve et al., 2010), although a decrease in the thickness of the INL and an increase in the thickness of the plexiform layers were observed in adults. This suggests that the increase in retinal size during postnatal development might not be mainly due to addition of new cells but rather to morphogenetic movements such as tissue narrowing by cell intercalation and extension and also to an increase in the extension/number of neuronal processes in plexiform layers.

Based in data obtained from teleosts (Johns, 1977; Julian et al., 1998; Marcus et al., 1999), it has been largely assumed that the retina of adult fishes presents active cell proliferation and neurogenesis (reviewed by Alunni and Bally-Cuif, 2016). However, our data shows that the catshark retina has virtually no mitotic activity after sexual maturation. Lack of cell proliferation has been also observed in the adult lamprey retina (Villar-Cheda et al., 2008), which suggests that the loss of active neurogenesis after sexual maturation is the ancestral character in vertebrates.

In the peripheral retina (CMZ and TZ), we also observed significant changes in the expression of progenitor markers. GFAP+ and *ScSox2*+ progenitors become highly restricted to the retinal margin in adults. Since the vast majority of these cells do not express PCNA or pH3 at this period, they could correspond to quiescent progenitors that keep their ability to re-enter the cell cycle in case of injury or disease, as previously described in teleost fish and amphibians (Fausett and Goldman, 2006; Langhe et al., 2017). Future work should determine whether mitotic activity, proliferation and neurogenesis are reactivated after injury in adult catsharks.

Finally, RNA-Seq analyses of the retinal transcriptome of juvenile and adult catsharks revealed a decrease in the expression of genes typically associated with proliferation, neurogenesis, cell-cycle regulation, and maintenance of progenitor cell properties. This confirmed that the catshark retina losses most of its proliferative and neurogenic activity after sexual maturation and that these genes probably play key roles in maintaining a high neurogenic activity in juveniles. We also observed changes in the expression of genes belonging to pathways known to be involved in the control of neurogenic processes like Shh, Wnt, Notch or Slit/Robo signaling (Supplementary Table 2). The RNA-Seq results fit perfectly with the histological and immunofluorescence data. Taken together, these data and those reported in other vertebrates show that the maintenance of an active proliferating state in the adult retina is a variable trait of vertebrates, suggesting a high evolvability of underlying mechanisms. Our RNA-Seq data provide an excellent resource to identify new genes and signaling pathways controlling neurogenesis in the vertebrate retina, to decipher the molecular basis for such variations and clarify underlying evolutionary modalities.

---

**Supplementary File 1**. Differentially expressed genes between juvenile and adult retinas.

**Supplementary File 2**. Gene Ontology (GO) and Kyoto Encyclopaedia of Genes and Genomes (KEGG) pathway enrichment analyses. Abbreviations: BP: biological process; CC: cellular component; MF: molecular function.

## 6. Author Contributions

All authors had full access to all the data in the study and take responsibility for the integrity of the data and the accuracy of the data analysis. IH-N, FA, AB-I, EC conceived and devised the study and approved the article. Study concept and design: IH-N, FA, AB-I, EC. Acquisition of data: IH-N, SM, LS, HM, DR. Analysis and interpretation of data: IH-N, DR, FA, AB-I, EC. Drafting of the manuscript: IH-N, DR, FA, AB-I, EC. Critical revision of the manuscript: IH-N, SM, LS, HM, DR, FA, AB-I, EC. Obtained funding: IH-N, EC.

