## Supplemental Table 1 for "Loss of active neurogenesis in the adult shark retina"

**Supplementary Table 1**. Primary and secondary antibodies used in double immunofluorescence experiments.

| **PRIMARY ANTIBODY** | **DILUTION** | **REFERENCE** | **SECONDARY ANTIBODY** | **DILUTION** | **REFERENCE** |
| --- | --- | --- | --- | --- | --- |
| **Rabbit polyclonal anti-GFAP** | 1:500 | DAKO Z033429 | Cy3-goat anti-rabbit IgG (H+L) | 1:200 | Invitrogen A10520 |
| **Mouse monoclonal anti-GS** | 1:500 | Millipore Mab302 | FITC-goat anti-mouse IgG (H+L) | 1:200 | Invitrogen F2761 |
| **Rabbit polyclonal anti-pH3** | 1:300 | Millipore  06-570 | Cy3-goat anti-rabbit IgG (H+L) | 1:200 | Invitrogen A10520 |
| **Mouse monoclonal anti-PCNA** | 1:500 | Sigma P8825 | FITC-goat anti-mouse IgG (H+L) | 1:200 | Invitrogen F2761 |
