## Supplemental Table 2 for "Loss of active neurogenesis in the adult shark retina"

**Supplementary Table 2**. Genes belonging to developmental signalling pathways that showed differential expression in the RNA-Seq analyses. References linking these signalling pathways to retinal development are indicated in the third column.

| **PATHWAY** | **GENES** | **REFERENCES** |
| --- | --- | --- |
| **Shh pathway** | GLI2  (Zinc finger protein GLI2) | Wall et al., 2001; Sanek et al., 2009; Todd and Fischer, 2015; Gallardo and Bovolenta, 2018 |
|  | HHIP  (Hedgehog-interacting protein) |  |
|  | HES1B  (Transcription factor HES1B) |  |
|  | ZIC2  (Zinc finger protein 2) |  |
| **Wnt pathway** | WNT10A  (protein Wnt10a) | Hunter et al., 2004; Poggi et al., 2018 |
|  | WIF1  (Wnt inhibitory factor 1) |  |
|  | DKK3  (Dickkopf-related protein 3) |  |
|  | GBP  (GSK3-binding protein) |  |
| **Notch pathway** | NOTCH1  (Neurogenic locus notch  homolog protein 1) | Lindsell et al., 1996; Nelson et al., 2009; Maurer et al., 2014; Reisenberg and Brown, 2016; Ivanov, 2019; Lee et al., 2020 |
|  | DLK1  (Protein delta homolog 1) |  |
|  | MAML2  (Mastermind-like protein 2) |  |
|  | JAG2  (Jagged-2) |  |
|  | HES1B  (Transcription factor HES 1B) |  |
| **Slit-Robo pathway** | ROBO1  (Roundabout homolog 1) | Erskine et al., 2000; Huang et al., 2009 |
|  | ROBO2  (Roundabout homolog 2) |  |
|  | SLIT3  (Slit homolog 3 protein |  |
|  | SRGP2  (SLIT-ROBO Rho GTPase-activating protein 2) |  |
|  | SRGP3  (SLIT-ROBO Rho GTPase-activating protein 3) |  |
|  | MYCN  (N-myc proto-oncogene protein) | Xue and Harris, 2011 |
|  | MYBP  (C-myc binding protein) |  |
|  | NMI  (N-myc interactor) |  |
